# Distinct roles for innexin gap junctions and hemichannels in mechanosensation

**DOI:** 10.1101/716324

**Authors:** Denise S. Walker, William R. Schafer

## Abstract

Mechanosensation is central to a wide range of functions, including tactile and pain perception, hearing, proprioception, and control of blood pressure, but identifying the molecules underlying mechanotransduction has proved challenging. In *Caenorhabditis elegans*, the avoidance response to gentle body touch is mediated by 6 touch receptor neurons (TRNs), and is dependent on MEC-4, a DEG/ENaC channel. We show that hemichannels containing the innexin protein UNC-7 are also essential for gentle touch in the TRNs, as well as harsh touch in both the TRNs and the PVD nociceptors. UNC-7 and MEC-4 do not colocalize, suggesting that their roles in mechanosensory transduction are independent. Heterologous expression of *unc-7* in touch-insensitive chemosensory neurons confers ectopic touch sensitivity, indicating a direct role for UNC-7 hemichannels in mechanosensation. The *unc-7* touch defect can be rescued by the homologous mouse gene *Panx1* gene, thus, innexin/pannexin proteins may play broadly conserved roles in neuronal mechanotransduction.

## Introduction

Innexins are a family of proteins that form gap junctions in invertebrate neurons and muscle cells. Gap junctions allow free (though gated) movement of ions and small signalling molecules between the cytoplasm of the connected cells, resulting in electrical coupling and the propagation of signals such as Ca^2+^ waves. Like in vertebrates, where gap junctions are formed from unrelated proteins called connexins, each invertebrate gap junction consists of two innexin hemichannels, each of which is a hexamer of constituent subunits (Phelan and Starich, 2001). The innexin families can be relatively large; for example, *C. elegans*, where innexins were originally identified, has 25 innexin genes. Different family members have distinct expression patterns, distinct gating properties, and differ in their ability to form homo- or hetero-hexamers and homo- or heterotypic gap junctions with specific partner hemichannels. There is thus enormous potential for variety, as well as asymmetry (rectification) in the relationships between partner cells (Hall, 2019; Palacios-Prado et al., 2014; Phelan et al., 2008).

In addition to their roles in gap junctions, the constituent hemichannels can also function independently as gated channels connecting the cell’s cytoplasm with the exterior. Hemichannels have been shown to be gated by a variety of stimuli, including changes in extracellular pH, Ca^2+^ concentration, or mechanical stimulation (Herve and Derangeon, 2013; Saez et al., 2005). Indeed, the vertebrate homologues of innexins, the pannexins, (Baranova et al., 2004; Bruzzone et al., 2003) (Yen and Saier, 2007), are thought to function exclusively as channels (i.e. pannexons), not as gap junctions (Sosinsky et al., 2011). Humans have 3 pannexin genes, and it is becoming increasingly evident that they play an important role in a wide range of medically significant processes, such as apoptosis, inflammation, ischemia and tumour genesis (Chiu et al., 2014; MacVicar and Thompson, 2010; Penuela et al., 2013), as well as neuropathic pain (Jeon and Youn, 2015). Given the homology between innexins and pannexins, *C. elegans* represents an amenable system in which to gain a greater understanding of these “hemichannel” functions, as well as the role of gap junctions in the organisation and function of neuronal circuits.

Perhaps the best characterized innexin genes in *C. elegans* are *unc-7* and *unc-9*. Mutations in these genes were originally identified based on the uncoordinated locomotion phenotype caused by loss of UNC-7 or UNC-9 function (Brenner, 1974). Both genes are expressed in ventral cord motorneurons and the premotor interneurons that promote forward or backward crawling (Altun et al., 2009; Starich et al., 2009). Heterotypic gap junctions between these premotor interneurons and motorneurons are important for controlling the balance between forward and backward locomotion as well as promoting coordinated sinusoidal locomotion (Kawano et al., 2011; Starich et al., 2009). They also play a central role in the regulation of sleep (Huang et al., 2018). In addition, UNC-7 has been shown to function as a hemichannel in motorneurons to promote neuromuscular activity through regulation of presynaptic excitability (Bouhours et al., 2011). UNC-7 has also been shown to function in the sensory circuit involved in nose touch, most likely through gap junctions in a hub-and-spoke electrical circuit (Chatzigeorgiou and Schafer, 2011). Both UNC-7 and UNC-9 are expressed in many additional neurons, where their functions have not been investigated.

Among the cells that express *unc-7* and *unc-9* are the sensory neurons mediating gentle and harsh body touch. Six neurons (referred to as TRNs or gentle touch neurons) are involved in sensing gentle touch: the ventral AVM and PVM, and lateral pairs of ALMs and PLMs. A mechanosensory complex including the DEG/ENaC channel subunits MEC-4 and MEC-10 is required for gentle touch responses in all these neurons (Bianchi, 2007; Bounoutas and Chalfie, 2007; Schafer, 2015). The anterior touch neurons form a putative gap junction-coupled electrical network, with ALML and ALMR coupled to AVM, as well as to the locomotion circuit via AVDR (Chen et al., 2006; Hall and Russell, 1991; Varshney et al., 2011; White et al., 1986). In contrast, the posterior TRNs are not gap junction coupled, though they do make gap junctions with other neurons. The PVD neurons, which sense harsh body touch, also express both *unc-7* and *unc-9* but the extent to which they form gap junctions is unclear. While gap junctions were not detected previously (Varshney et al., 2011; White et al., 1986), this may be due to their complex, branched morphology; more recent analysis (Cook et al., 2019) identified a few gap junctions with motorneurons. Since pannexin 1 has been demonstrated to function as mechanosensitive, ATP releasing channels in multiple cellular contexts (Bao et al., 2004; Beckel et al., 2014; Furlow et al., 2015; Richter et al., 2014), this might suggest a role for innexin hemichannels in mechanotransduction in the PVDs and the TRNs.

In this study, we characterise the roles of two innexin subunits, UNC-7 and UNC-9, in *C. elegans* touch neurons. Both UNC-7 and UNC-9 are required for gap junction communication between the anterior TRNs, creating an electrically-coupled network that ensures a robust response to stimuli applied to either side of the animal. In addition, UNC-7 hemichannels play an essential role in gentle touch mechanosensation in both the anterior and posterior TRNs as well as harsh touch sensation in the PVD polymodal nociceptors. Heterologous expression of UNC-7 hemichannels in mechanically insensitive olfactory neurons confers the ability to respond to nose touch, indicating that UNC-7 is sufficient as well as necessary to generate a mechanosensor. Since mouse pannexin 1 can functionally complement an *unc-7* null mutation, our results may suggest conserved functions of pannexins and innexins in other mechanosensory tissues.

## Results

### Innexins are required for mechanosensation and electrical coupling of touch neurons

At least three innexin genes – *inx-7, unc-7* and *unc-9 –* have been shown to be expressed in the TRNs (Altun et al., 2009; Starich et al., 2009). We therefore used RNAi to investigate the role of these genes in mechanosensory activity. Since global knockdown could potentially have complex consequences, we used the touch neuron-specific *Pmec-7* promoter to cell-specifically inactivate each innexin gene and imaged neuronal touch responses using a genetically encoded Ca^2+^ indicator (Kerr et al., 2000; Suzuki et al., 2003). We observed that while knockdown of *inx-7* had no effect, knockdown of *unc-9* significantly reduced, and knockdown of *unc-7* almost completely abolished, ALM touch responses (**Figure 1B-D**). Thus, *unc-7* and *unc-9* both function in the gentle touch response in the TRNs.

**Figure 1.**
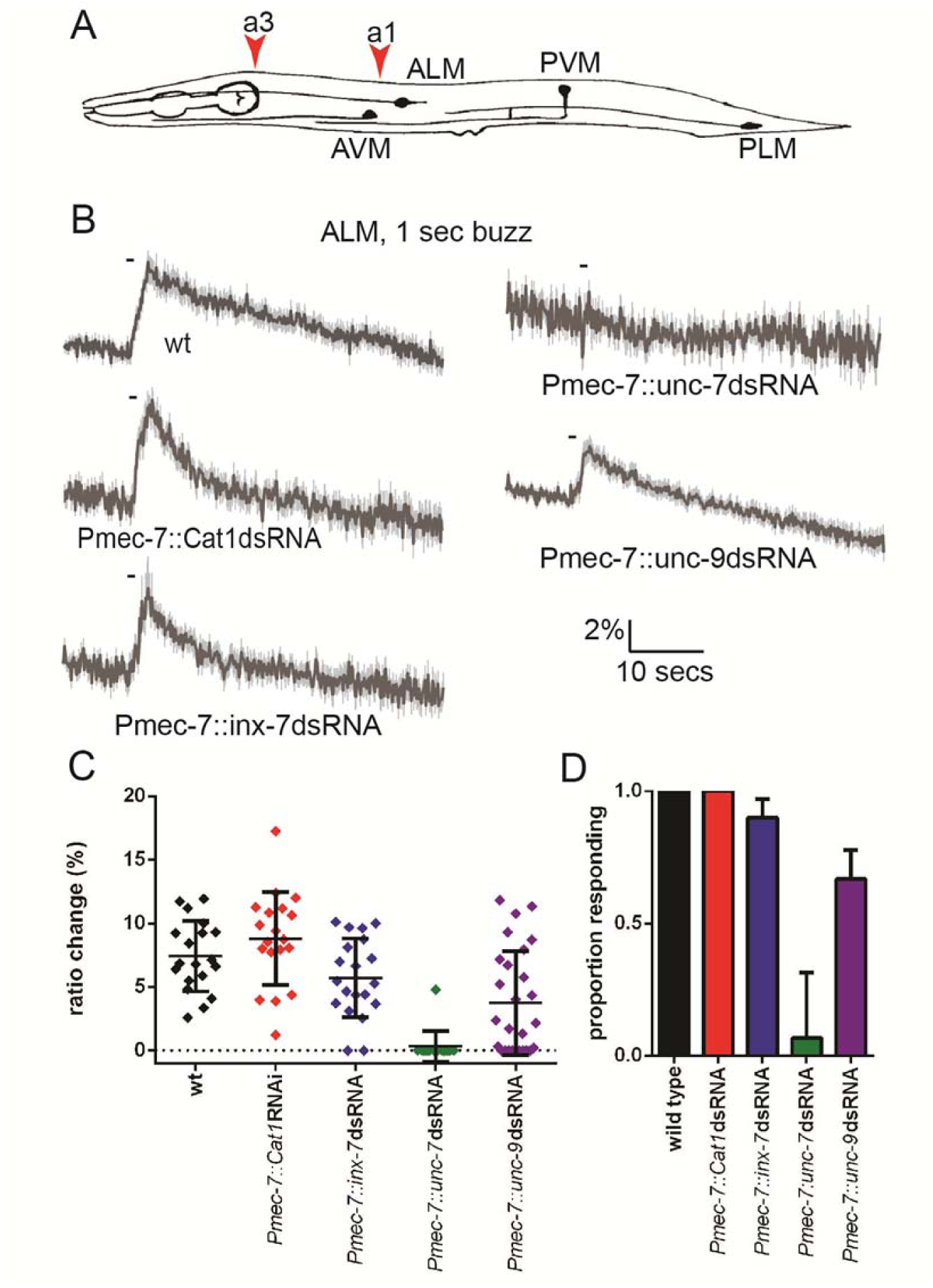
*unc-7* and *unc-9* function in touch responses in ALM. **(A)** Schematic showing positions of cell bodies and processes of the *C. elegans* touch receptor neurons. ALM and PLM are lateral pairs (left and right), of which only one of each is shown. Red arrowheads show stimulation sites. Except where stated, animals were stimulated at a3. **(B**,**C**,**D)** Gentle touch responses recorded in ALM for wild type animals and animals expressing dsRNA under control of *Pmec-7*. **(B)** Average traces of % ratio change. Grey indicates SEM. **(C)** Scatter plot showing individual ratio changes (diamonds). Bars indicate mean ± SEM. **(D)** Graph showing proportion responding. Error bars indicate SE. *unc-7* (<0.001) and *unc-9* (P<0.01) RNAi are significantly different from wild type, while *E. coli* Cat1 and *inx-7* RNAi are not, Fisher’s exact test).

The anterior touch receptor neurons are electrically-coupled through ALMR-AVM and ALML-AVM gap junctions; thus, these gap junctions could potentially influence touch responses. To investigate the importance of these electrical synapses in touch neuron activity, we first examined the consequences of laser ablating AVM, which would disrupt gap junction communication between ALML and ALMR. In wild type unablated animals, ALM responded robustly to ipsilateral (i.e. the left side for ALML) or contralateral (i.e. the right side for ALML) stimuli (**Figure 2A-C**). However, when AVM was ablated, ALM responded robustly only to ipsilateral stimuli. This indicates that electrical coupling between the anterior TRNs, via AVM, is required for touch neurons to respond to contralateral stimuli.

**Figure 2.**
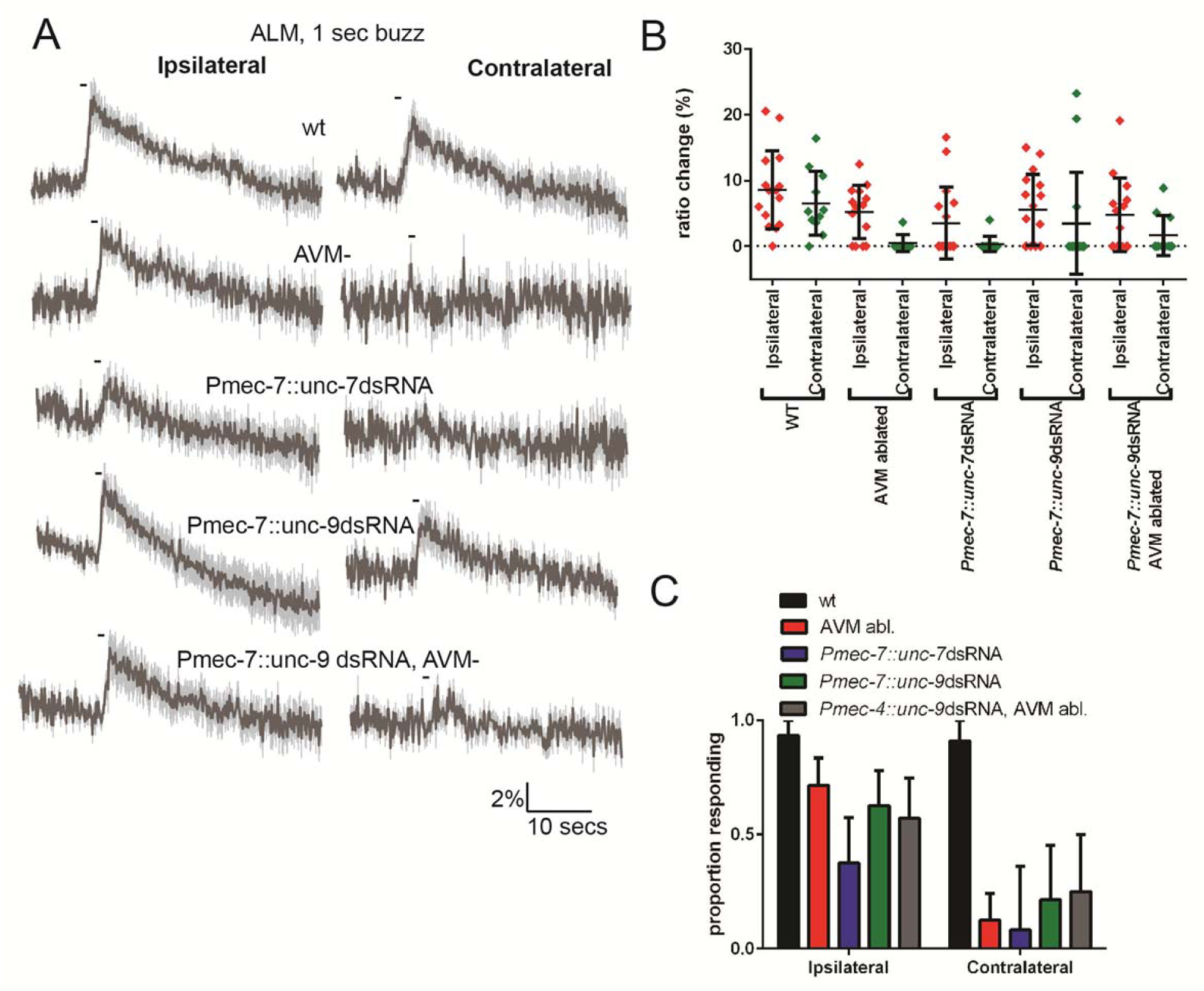
Innexins are required for mechanosensation and electrical coupling of touch neurons. Gentle touch responses recorded in ALM for wild type worms, worms in which AVM has been laser ablated and worms expressing dsRNA under control of *Pmec-7*. Neurons have been classified as “ipsilateral” or “contralateral”, according to the position of the cell body relative to the stimulation site and the hypothetical midline of the animal. **(A)** Average traces of % ratio change. Grey indicates SEM. **(B)** Scatter plot showing individual ratio changes (diamonds). Bars indicate mean ± SEM. **(C)** Graph showing proportion responding. Error bars indicate SE. In ipsilateral neurons, the proportion of AVM ablated animals or *unc-9* RNAi animals responding is not significantly different to wild type, while *unc-7* RNAi animals respond at a significantly reduced rate (P<0.01). Combining *unc-9* RNAi with AVM ablation is not significantly different from either *unc-9* RNAi alone or AVM ablation alone. In contralateral neurons, the proportions of AVM ablated, *unc-7* RNAi and *unc-9* RNAi animals responding are all significantly lower than wild type (P<0.01; P<0.001; P<0.01). Combining *unc-9* RNAi with AVM ablation is not significantly different from either *unc-9* RNAi alone or AVM ablation alone, Fisher’s exact test.

**Figure 2 – figure supplement 1.**
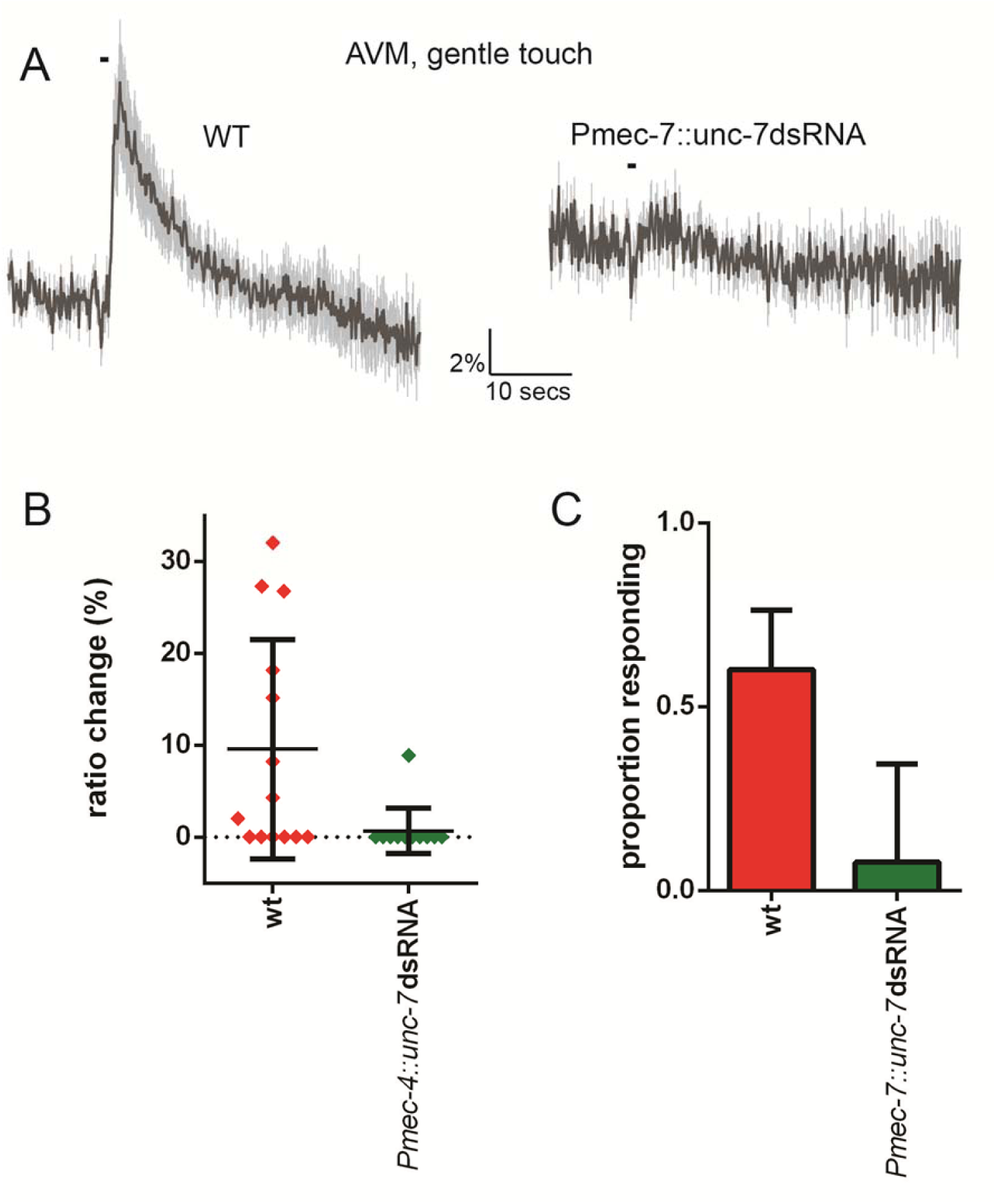
*unc-7* is required for gentle touch response in AVM. Ca^2+^ response to 1 sec buzz in AVM, for wild type and TRN-specific *unc-7* RNAi animals. All AVM neurons assayed were located in the “near” half of the animal, with respect to the stimulation site, as determined by the position of the cell body. **(A)** Average traces of % ratio change. Grey indicates SEM. **(B)** Scatter plot showing individual ratio changes (diamonds). Bars indicate mean ± SEM. **(C)** Graph showing proportion responding. Error bars indicate SE. The proportion of AVM “near” neurons responding is significantly lower in *unc-7* RNAi animals compared to wild type (P<0.05, Fisher’s exact test).

**Figure 2 – figure supplement 2.**
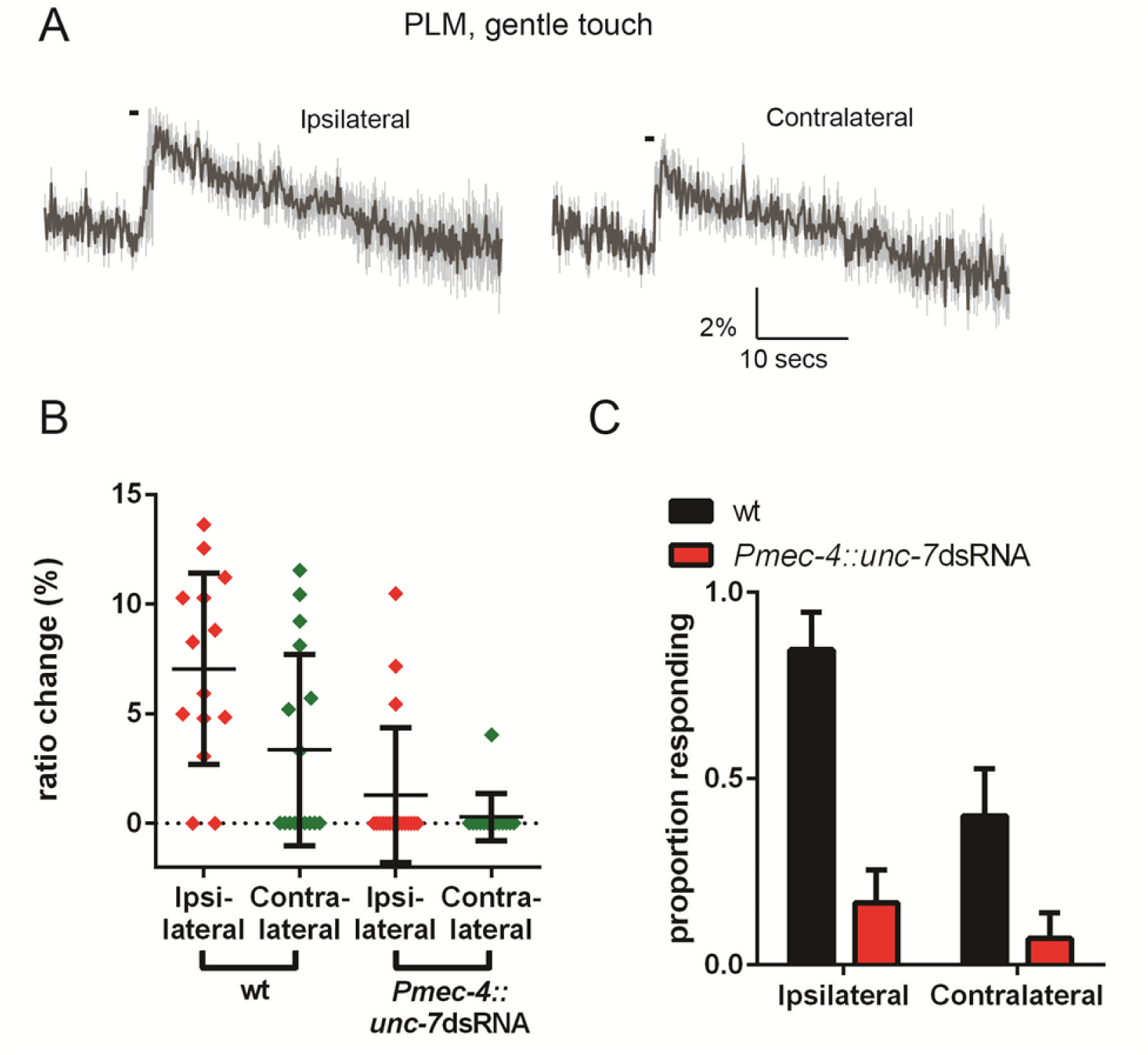
PLML and PLMR do not cooperate via gap junctions. **(A)** Average traces of % ratio change. Grey indicates SEM. **(B)** Scatter plot showing individual ratio changes (diamonds). Bars indicate mean ± SEM. **(C)** Graph showing proportion responding. Error bars indicate SEM. The proportion responding is significantly lower in “contralateral” neurons compared to those ipsilateral to the stimulation site (P<0.05, Fisher’s exact test).

To assess the possible roles of *unc-7* and *unc-9* in this electrical coupling, we re-examined the effects of innexin RNAi, distinguishing between ALM responses to ipsilateral and contralateral stimuli (**Figure 2A-C**). We observed that knockdown of *unc-9* almost entirely abolished responses to contralateral stimuli but had little effect on responses to ipsilateral stimuli, suggesting that UNC-9 is an important constituent of the gap junctions coupling the ALMs through AVM. Consistent with this possibility, the combined effect of *unc-9* RNAi and AVM ablation was not significantly different from either *unc-9* RNAi alone or AVM ablation alone in either ipsilateral or contralateral neurons (**Figure 2A-C**). Thus, disrupting gap junction communication appears functionally analogous to disrupting *unc-9*, supporting the hypothesis that *unc-9* plays an essential role in gap junction communication between the TRNs, and that this is probably its sole function in the TRNs. In contrast, *unc-7* RNAi significantly disrupted the responses in ALM to both ipsilateral and contralateral stimuli, suggesting that UNC-7 is required for mechanosensation *per se*, rather than simply contributing to gap junctions. As **Figure 2 – figure supplement 1** shows, *unc-7* RNAi also severely disrupts responses in AVM, even when the stimulus was applied close to the AVM dendrite. Together, these results indicate that *unc-7* is required for robust mechanosensory responses in all three anterior TRNs.

### The mechanosensory function of UNC-7 is gap junction-independent

In principle, the mechanosensory defects seen in the anterior touch receptor neurons could result from changes in excitability due to a lack of UNC-7-containing gap junctions; alternatively, UNC-7-containing hemichannels could have a distinct, gap-junction-independent role in mechanosensation. To address these possibilities, we first investigated touch responses in the posterior TRNs PLML and PLMR, which are not connected by gap junctions, either directly or indirectly via PVM. In wild type animals, PLM responses to ipsilateral stimuli were extremely robust (nearly 100% responding), while less than half of contralateral stimuli generated responses (**Figure 2 – figure supplement 2**), suggesting that the coordinated responses of the anterior TRNs are indeed gap-junction-dependent. When we measured responses in *unc-7* RNAi animals, we observed defective responses to both ipsilateral and contralateral stimuli, as seen previously for the anterior TRNs (**Figure 2 – figure supplement 2**). Thus, *unc-7* is appears to be required for the response in the posterior TRNs, despite their lack of gap junction interconnectivity.

Formation of gap junctions by innexins has been shown (Bouhours et al., 2011) to require four cysteines at the inter-hemichannel interface of UNC-7. When these residues are mutated, the innexin protein’s ability to form functional gap junctions is disrupted, but its hemichannel function is intact. We therefore examined whether a “cysless” mutant allele of *unc-7* could rescue the *unc-7* mechanosensory defect in touch neurons. As **Figure 3A-C** shows, ALM gentle touch responses are severely disrupted in *unc-7(e5)* animals, as we observed previously for *unc-7* RNAi. Expression of a wild type *unc-7* cDNA (isoform a, also known as UNC-7L (Starich et al., 2009)) under the control of the TRN-specific *mec-4* promoter significantly rescued this defect. Expression of a cysless mutant cDNA also very successfully rescued, indicating that the TRN defect is related to hemichannel rather than gap junction activity. Expression of mouse *Panx1* (encoding *Pannexin 1*), but not *Panx2*, in the TRNs also successfully rescued the *unc-7* mechanosensory defect (**Figure 3A-C**), indicating that despite the relatively low sequence similarity, UNC-7 and Pannexin 1 share significant functional conservation. Intriguingly, expression of a cDNA encoding a shorter *unc-7* isoform (“c” or UNC-7SR (Starich et al., 2009)) which is known to form gap junctions failed to rescue (**Figure 3A-C**). Thus, UNC-7’s mechanosensory function appears to be genetically-separable from its ability to form gap junctions, and may therefore specifically involve hemichannels.

**Figure 3.**
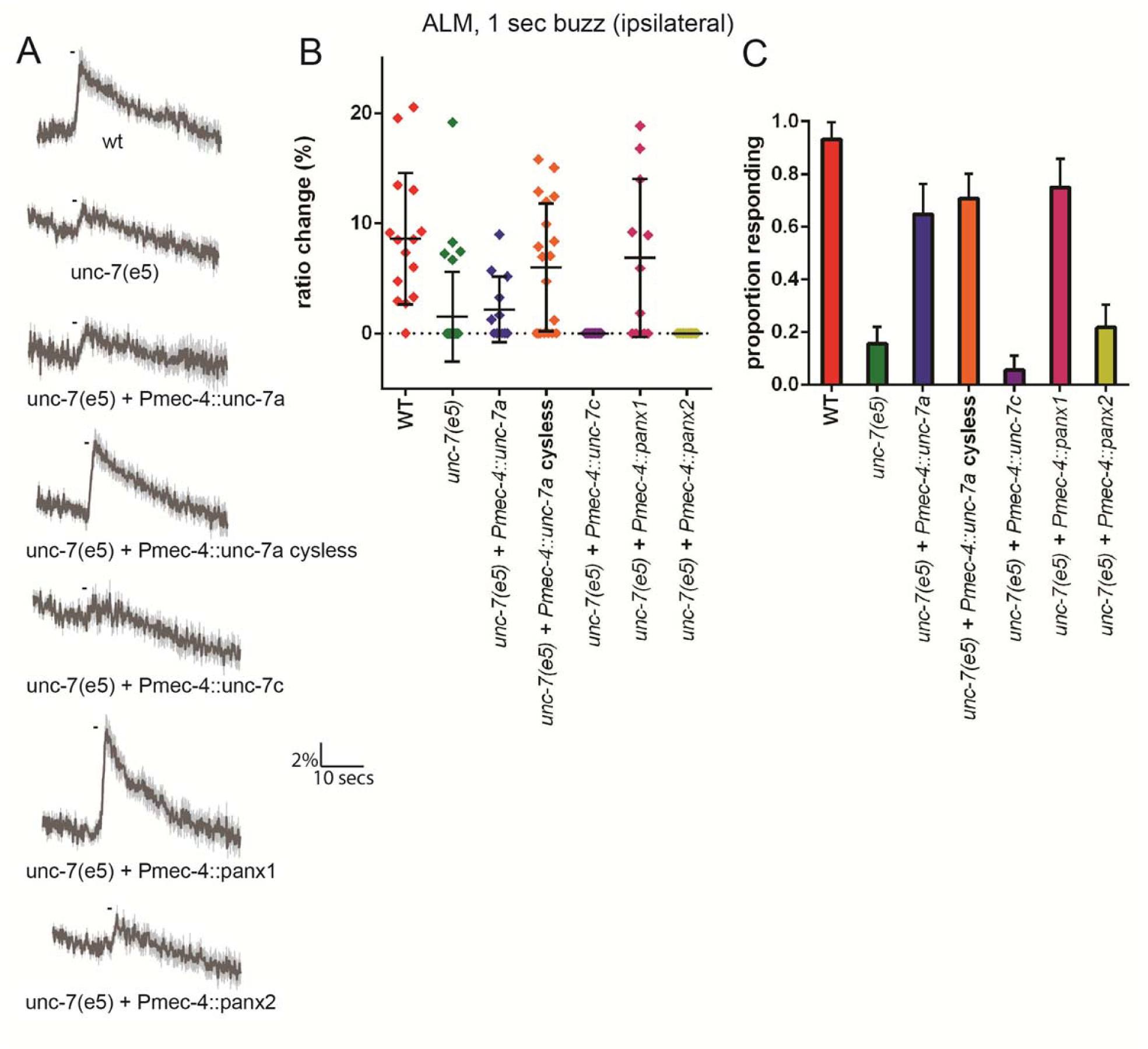
The mechanosensory function of UNC-7 is gap junction-independent. Gentle touch responses recorded in ALM for wild type, *unc-7(e5)*, and *unc-7(e5)* animals expressing *unc-7* isoforms or pannexins under the control of *Pmec-4*. “Cysless” indicates C173A, C191A, C377A, C394A. All neurons recorded were ipsilateral, according to the position of the cell body relative to the stimulation site and the hypothetical midline of the animal. **(A)** Average traces of % ratio change. Grey indicates SEM. **(B)** Scatter plot showing individual ratio changes (diamonds). Bars indicate mean ± SEM. **(C)** Graph showing proportion responding. Error bars indicate SE. The response frequency is significantly reduced in *unc-7*(e5) compared to wildtype (P<0.001). This is significantly rescued by TRN expression of wild type (P<0.01) or cysless *unc-7a* (P<0.001). Cysless *unc-7* still significantly rescued the mutant when AVM was ablated (P<0.001), and there was no significant difference between AVM ablated and unablated cysless *unc-7*-expressing animals. While *unc-7c* and mouse *panx2* did not significantly rescue, *panx1* did (P<0.001), Fisher’s exact test).

### *unc-7* is specifically required for mechanosensation in touch neurons and nociceptors

In principle, UNC-7 could affect mechanosensory responses by affecting the excitability of the touch neurons; alternatively, UNC-7 could play a direct role in mechanosensation. To address these possibilities, we examined the effect of *unc-7* knockdown and overexpression on channelrhodopsin-mediated activation of the TRNs. When channelrhodopsin is expressed in the TRNs, photostimulation evokes an escape response similar to those evoked by mechanosensory stimulation (Nagel et al., 2005). To assess whether *unc-7* RNAi affects TRN excitability, we chose a stimulus duration at which only two thirds of wild type animals responded. As expected, a *mec-4* null mutation did not significantly alter the proportion of animals responding, consistent with the specific role played by *mec-4* in mechanotransduction. Likewise, neither *unc-7* RNAi nor overexpression of “cysless” *unc-7* significantly altered the proportion of animals responding to light stimulation (**Figure 4A**) suggesting that *unc-7* also does not alter the excitability of the touch neurons. The basal calcium activity of the touch neurons, as indicated by the baseline YFP/CFP ratio, also showed no significant difference between wild type animals (2.01±0.22) and *unc-7(e5)* (1.82±0.32). Together, these results indicate that loss of UNC-7 does not affect touch neuron excitability, and its role is likely to be specific to mechanosensation.

**Figure 4.**
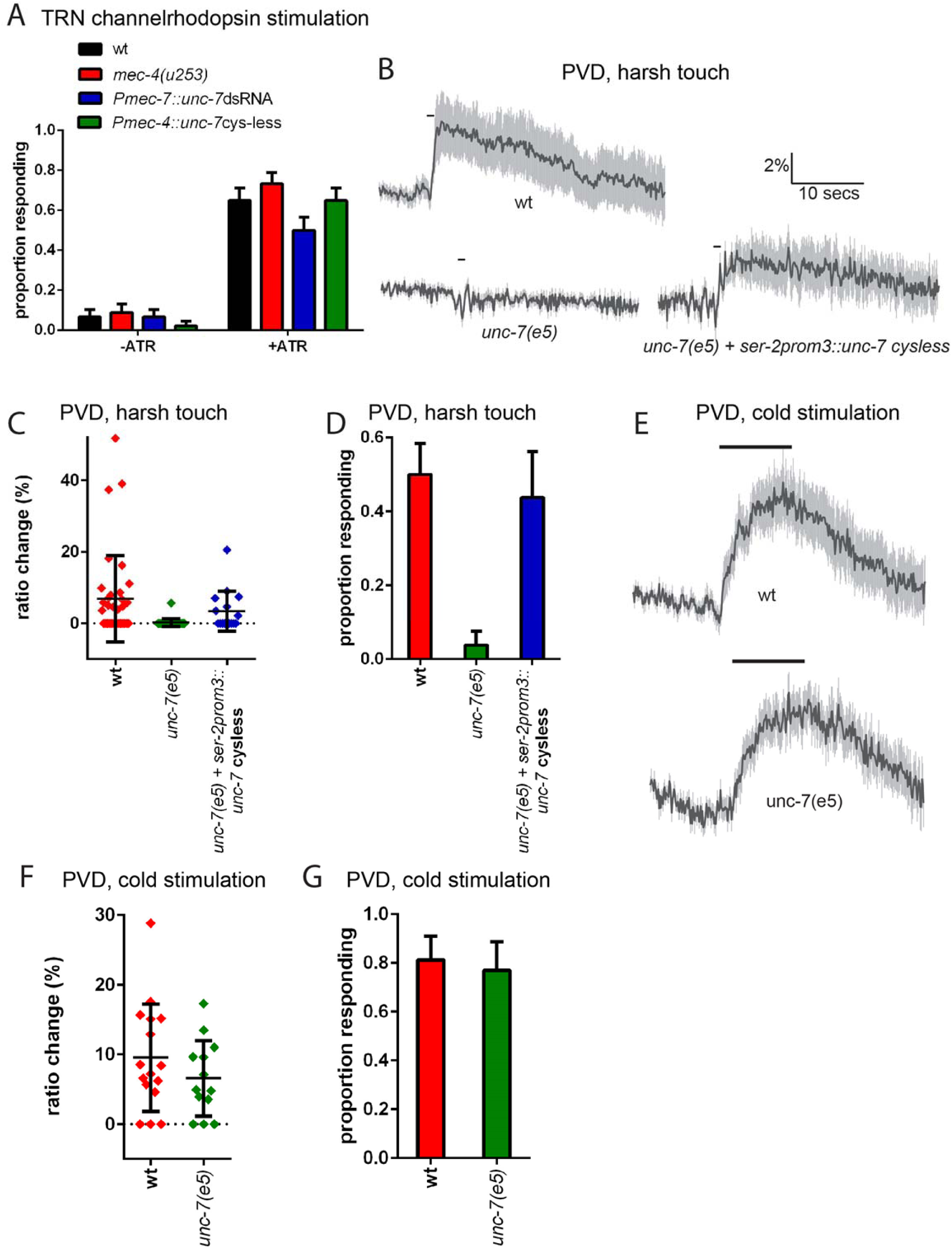
*unc-7* is specifically required for mechanosensation. **(A)** Behavioural response to light stimulation of animals expressing channelrhodopsin in the TRNs. The proportion of wild type animals responding was not significantly different to that for *Pmec-7∷unc-7*dsRNA, *Pmec-4∷unc-7* cysless or *mec-4(u253)* animals. **(B**,**C**,**D)** Harsh touch responses recorded in PVD for wild type and *unc-7(e5)* animals. **(B)** Average traces of % ratio change. Grey indicates SEM. **(C)** Scatter plot showing individual ratio changes (diamonds). Bars indicate mean ± SEM. **(D)** Graph showing proportion responding. Error bars indicate SE. The proportion responding is significantly reduced in *unc-7(e5)* animals (P<0.001); and this is significantly rescued (P<0.01) to a response rate not significantly different from wild type. **(E**,**F**,**G)** Cold responses recorded in PVD for wild type and *unc-7(e5)* animals. **(E)** Average traces of % ratio change. Black bar indicates shift from 22°C to 15°C. Grey indicates SEM. Scatter plot showing individual ratio changes (diamonds). Bars indicate mean ± SEM. Graph showing proportion responding. Error bars indicate SE. The proportion responding is not significantly different, Fisher’s exact test).

*unc-7* is expressed in other sensory neurons, including the polymodal nociceptor PVD. PVD neurons respond to several aversive stimuli, including harsh touch and cold temperature (Chatzigeorgiou et al., 2010). To examine whether *unc-7* functions specifically in mechanosensation, we assayed the effect of *unc-7* mutations on both thermal and mechanical responses in PVD. We observed (**Figure 4B-D**) that *unc-7(e5)* animals were severely defective in the Ca^2+^ response of the PVD neurons to harsh touch. In contrast (**Figure 4E-G**), *unc-7(e5)* animals showed no significant difference compared to wild-type in the PVD response to cold (temperature shift from 22° to 15°). Thus, *unc-7* is required for mechanosensory responses, but dispensable for thermosensory responses, in PVD, suggesting a specific role for UNC-7 in mechanotransduction.

The TRNs also exhibit responses to harsh touch that are distinct from those to gentle touch. While gentle touch responses are transient, low amplitude, and seen in response to a wide range of stimulus speed and displacement, harsh touch responses are longer, higher amplitude and seen only in response to fast, high-displacement stimuli [Suzuki 2003; **Figure 5A**). In response to a harsh stimulus, wild type animals exhibit a mixture of these response types and, consistent with earlier work (Suzuki et al., 2003), we observed that the harsh-like response was not disrupted by mutations in *mec-4* (**Figure 5B, C**). In contrast, we found that *unc-7* knockdown by RNAi specifically disrupted the harsh-like response, while *unc-7(e5)* mutations eliminated virtually all TRN response to harsh touch. Expression of cysless *unc-7* in the TRNs significantly rescued the harsh touch response defect in *unc-7* mutant animals, indicating that the harsh touch defect in the mutant was at least partially due to the cell-autonomous mechanosensory activity of UNC-7 hemichannels. *unc-7* knockdown in combination with *mec-4* null abolished both types of response, suggesting that UNC-7 and MEC-4 act independently in the touch neurons.

**Figure 5.**
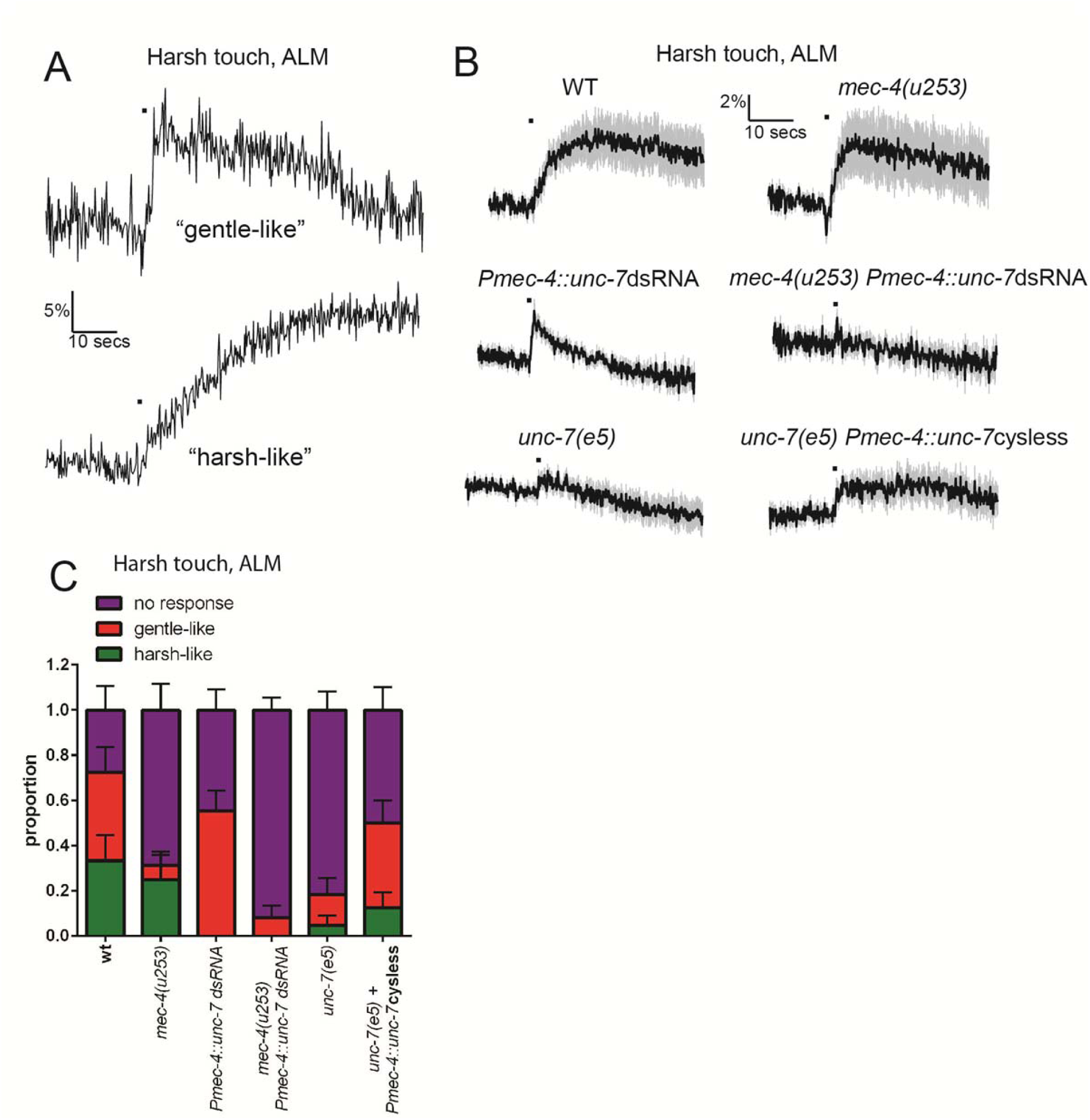
UNC-7 and MEC-4 have distinct roles in harsh touch. Ca^2+^ responses recorded in ALM in response to harsh touch. **(A)** Representative examples of the two types of Ca^2+^ responses to harsh touch stimulation in ALM. **(B)** Average traces of % ratio change. Grey indicates SEM. **(C)** Proportion of animals displaying the indicated types of Ca^2+^ response to harsh touch in ALM. Error bars are SE. The gentle-like response type is significantly disrupted in the absence of *mec-4* (P<0.05), while the harsh-like response is significantly disrupted by *unc-7* knockdown (P<0.01). *unc-7* RNAi, *mec-4* null combined completely abolishes both types of response (N.S. when compared to zero responses; P<0.001 when total response rate is compared to *unc-7* RNAi; N.S. when compared to mec-4 null). *unc-7(e5)* significantly disrupts the total response rate (P<0.01) and this is significantly rescued by TRN expression of *unc-7 cysless* (P<0.05), Fishers exact test.

### UNC-7 and MEC-4 act independently in touch neuron mechanosensation

To investigate the relationship between UNC-7 and MEC-4 in the touch neurons, we used fluorescently tagged transgenes to compare their intracellular localization patterns. As described previously (Zhang et al., 2004), mCherry-tagged MEC-4 protein was distributed in a punctate pattern along the ALM and PLM dendrites (**Figure 6A**). GFP-tagged UNC-7 was also expressed in a punctate pattern in both touch receptor neuron types. However, little overlap was observed between UNC-7 and MEC-4 puncta in either cell type (**Figure 6A, B**). Thus, UNC-7 does not appear to physically associate with MEC-4-containing mechanotransduction complexes, consistent with a distinct functional role in touch sensation.

**Figure 6.**
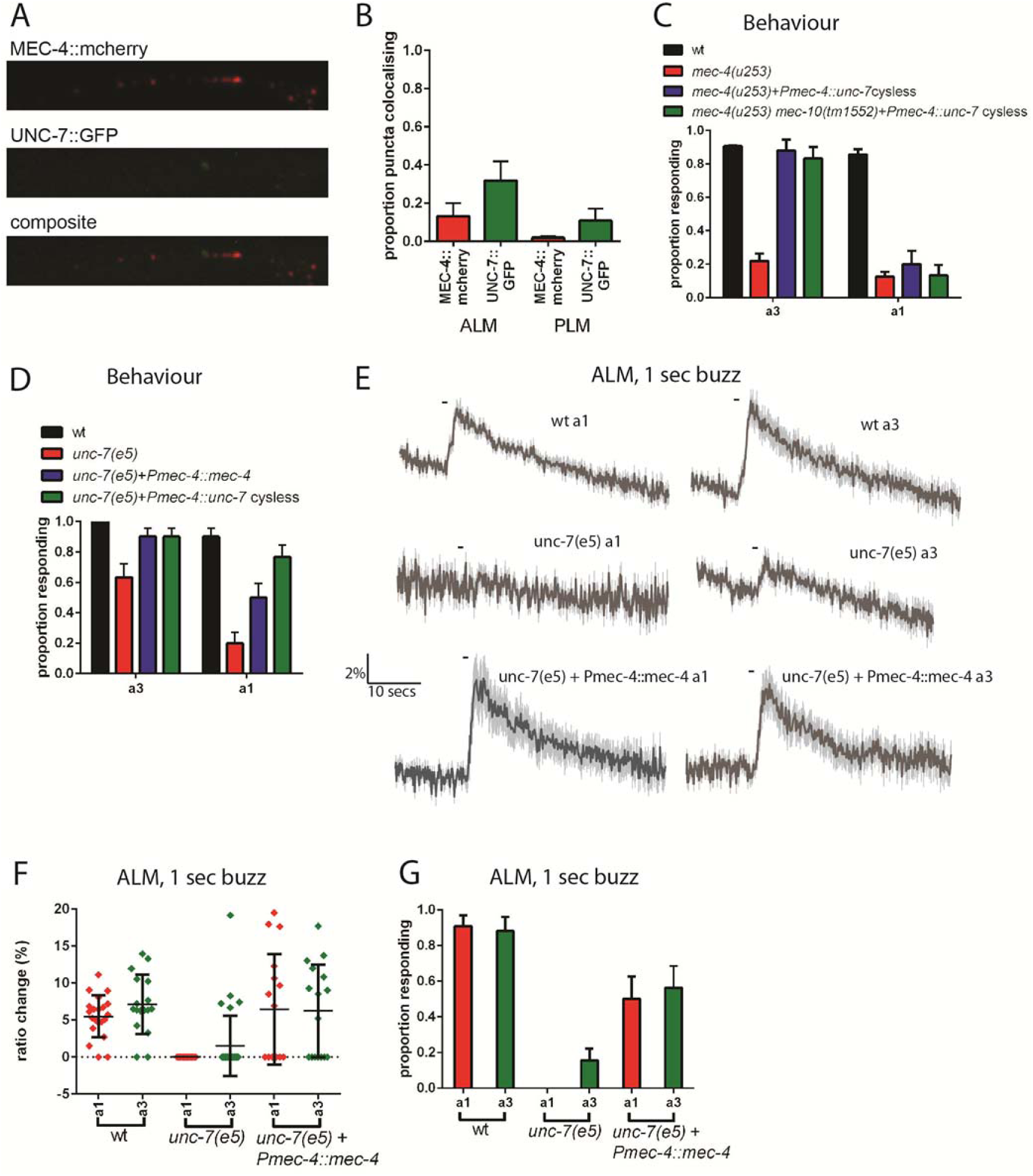
UNC-7 and MEC-4 act independently in touch neuron mechanosensation. **(A, B)** Confocal microscopy of TRN neurons expressing *mec-4*∷mcherry and *unc-7a*∷gfp. **(A)** Example images of PLM, and composite of the two channels, showing colocalisation in white. **(B)** Percentage of particles colocalising with particles of the other colour, based on centres of mass coincidence. **(C, D)** Behavioural response to anterior gentle touch, for genotypes indicated. Animals were stimulated either at the back of the terminal bulb (a3) or approximately 50µm anterior of the cell body of ALM (a1). Error bars are SE. TRN expression of *unc-7* cysless significantly rescued the behavioural defect of *mec-4(u253)*, including when *mec-10* was also defective, when stimulated at a3 (P<0.001 for both); but not when stimulated at a1. *unc-7(e5)* animals are significantly defective in the behavioural response to gentle touch at a3 (P<0.001) and a1 (P<0.001), and TRN expression of *mec-4* significantly rescued this, at a3 (P<0.05) and a1 (P<0.05). **(E**,**F**,**G)** Ca^2+^ responses to gentle touch recorded in ALM, for wild type, *unc-7(e5)*, and *unc-7(e5)* animals expressing P*mec-4∷mec-4.* **(E)** Average traces of % ratio change. Light grey indicates SEM. **(F)** Scatter plot showing individual ratio changes (diamonds). Bars indicate mean ± SEM. **(G)** Graph showing proportion responding. Error bars indicate SE. Expression of *mec-4* significantly rescued the Ca^2+^ response defect of *unc-7(e5)*, whether stimulated at a3 or a1 (P<0.01 for both), Fisher’s exact test.

Like *unc-7, mec-4* is critical for the response to gentle touch, and null mutations in *mec-4* result in an almost complete loss of touch-evoked Ca^2+^ response (Suzuki et al., 2003). We reasoned that if UNC-7 hemichannels act independently of MEC-4, then their overexpression in the TRNs might compensate for the absence of MEC-4. Indeed, when cysless *unc-7* was overexpressed in the TRNs (**Figure 6C**) we observed strong suppression of the *mec-4(u253)* defect in the behavioural response to gentle touch. This suppression was independent of *mec-10*, the other DEG/ENaC known to function in the TRNs. Interestingly, *unc-7* overexpression only restored touch responses to stimuli applied near the head (a3, **Figure 1A**), and we were unable to detect any rescue of Ca^2+^ responses in ALM regardless of the site of stimulation. This suggests that UNC-7 overexpression only partially compensates for loss of MEC-4, and is therefore insufficient to generate calcium transients in the ALM cell body, though it may be sufficient to activate local depolarization in presynaptic regions near the nerve ring. Conversely, we also tested whether overexpression of MEC-4 could compensate for the absence of UNC-7. We observed (**Figures 6D, E, F, G**) that when *mec-4* was overexpressed in the TRNs it strongly suppressed the behavioural and calcium defects in *unc-7(e5)* in response to either anterior or midbody touch stimuli. Thus, although both *mec-4* and *unc-7* are essential for the response to gentle touch in the TRNs, both can, at least to some extent, substitute for the other when overexpressed. This suggests that MEC-4 and UNC-7 indeed act independently as mechanotransducers in the TRNs.

### Heterologous expression of UNC-7 hemichannels in olfactory neurons confers touch sensitivity

Our results so far indicate that UNC-7 is necessary for normal mechanosensation in the TRNs. If UNC-7 hemichannnels play a direct role in mechanotransduction, they might also be expected to be sufficient to confer mechanosensory responses in cells that are natively touch-insensitive. To test this possibility, we expressed the cysless derivative of *unc-7* in the ASKs, a pair of ciliated chemosensory neurons that sense lysine and pheromones (Macosko et al., 2009; Wakabayashi et al., 2009), but do not respond to mechanical stimulation (**Figure 7**). We then assayed mechanosensory activity potentially conferred by the heterologously expressed transgene by measuring touch-evoked neural activity using an ASK-expressed genetically-encoded calcium indicator.

When we expressed the cysless *unc-7* transgene alone, we observed robust nose touch responses in ASK that were absent in the ASK neurons of wild-type animals (**Figure 7**). In contrast, expression of *mec-4* in the same way did not render ASK mechanically sensitive, even when coexpressed with *mec-2*. Coexpression of either *mec-2* or *mec-4* with cysless *unc-7* did not enhance the ectopic touch responses in ASK; indeed, the responses of coexpressing animals were if anything smaller than those of animals expressing *unc-7* alone. These results indicate that UNC-7 hemichannels specifically confer mechanical sensitivity in a heterologous cell type. Moreover, UNC-7 may require few if any additional specific factors to form a mechanosensor, whereas MEC-4 appears to require additional proteins to generate a mechanotransduction complex in the ASK neurons.

**Figure 7.**
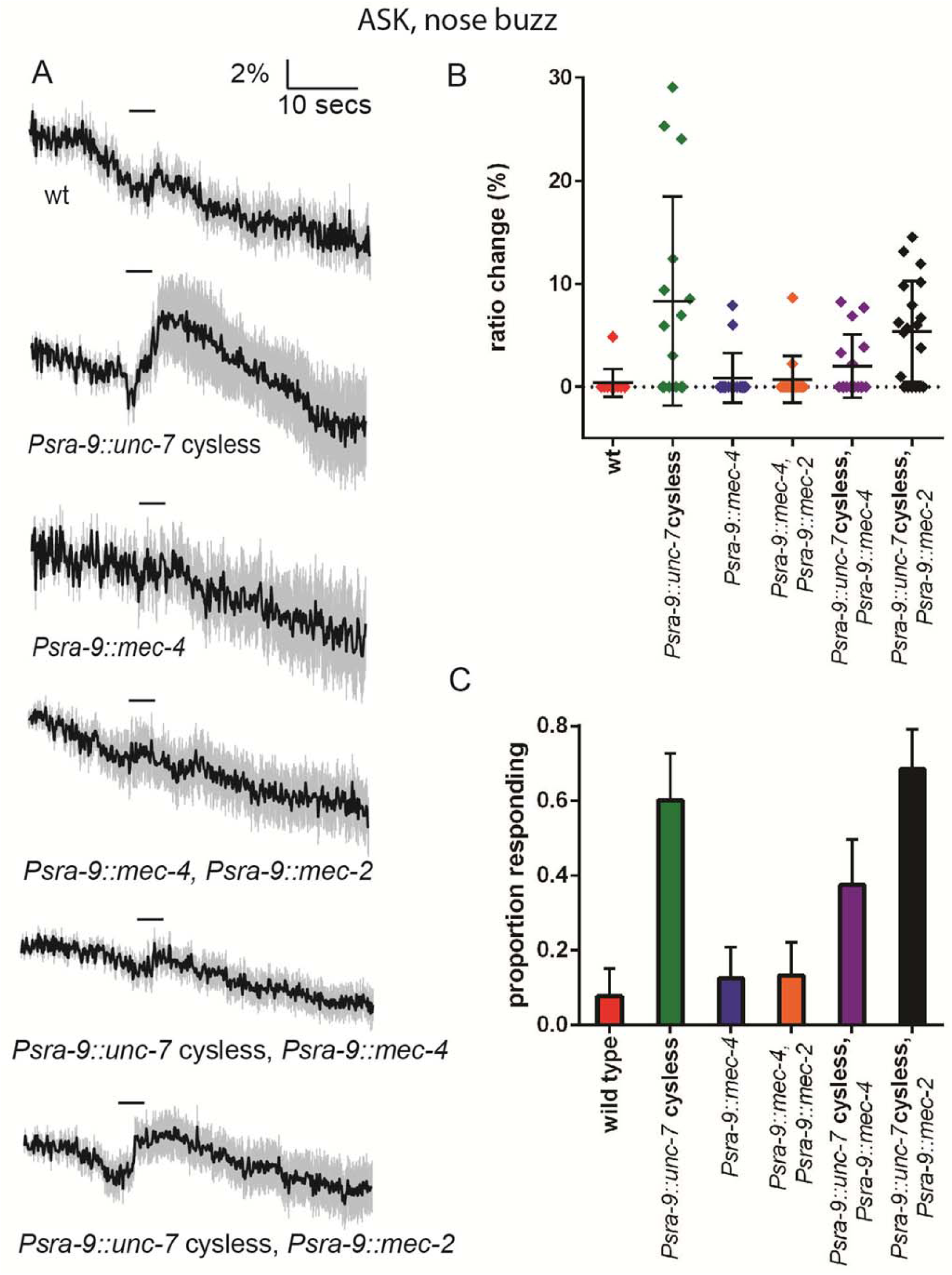
Heterologous expression of UNC-7 hemichannels in olfactory neurons confers touch sensitivity. Nose touch responses recorded in ASK of wild type animals and animals expressing *unc-7* cysless or *mec-4* in ASK. **(A)** Average traces of % ratio change. Light grey indicates SEM. **(B)** Scatter plot showing individual ratio changes. Bars indicate mean ± SEM. **(C)** Graph showing proportion responding. Error bars indicate SE. Wild type ASK neurons do not significantly respond to nose touch, but expression of *unc-7* cysless significantly increases the response rate (P<0.01). Expression of *mec-4* does not significantly increase the response rate, and coexpression of *mec-4* does not significantly alter the response rate for *unc-7* cysless expressing animals. Coexpression of *mec-2* does not significantly increase the response rate for *mec-4* or *unc-7*, Fisher’s exact test).

## Discussion

### UNC-7 hemichannels function specifically in mechanosensation

We have shown here that the innexin UNC-7 plays an essential role in the response to gentle touch, and that this mechanosensory function is likely mediated by hemichannels rather than gap junctions. Several lines of evidence support these conclusions. First, loss of *unc-7* function affects touch responses to ipsilateral stimuli as well as to contralateral stimuli, implying its role is not merely in indirect activation through gap junctions. Second, *unc-7* mutations lead to mechanosensory defects in neurons such as the PLMs, which are not known to be connected by gap junctions, and this may also be the case for the PVDs. Third, an *unc-7* transgene containing mutations that render it incapable of gap junction formation still effectively rescues the *unc-7* touch-insensitive phenotype. Our observation that *unc-7* functions affects mechanosensory, but not thermosensory responses in the PVD polymodal nociceptors, coupled with the fact that *unc-7* does not impair channelrhodopsin-mediated excitation of the TRNs, provide evidence that the UNC-7 hemichannels specifically affect mechanosensory transduction rather than neuronal excitability. Finally, heterologous expression of UNC-7 hemichannels in a chemosensory neuron conferred mechanosensitivity, indicating a specific and direct role in mechanotransduction.

How might UNC-7 contribute to mechanotransduction in the touch neurons and PVD nociceptors? Perhaps the simplest hypothesis is that UNC-7 hemichannels are themselves mechanically-gated ion channels whose opening contributes to the mechanoreceptor potential. Consistent with this possibility, pannexin 1, a mammalian homologue of UNC-7 that functionally complements its touch phenotype in worms, has been shown to form mechanosensitive channels when expressed in Xenopus oocytes (Bao 2004). However, although heterologously-expressed UNC-7 (in the cysless mutant form) appears to have channel activity, the potential mechanosensitivity of these channels was not reported (Bonhours 2011). Alternatively, UNC-7 hemichannels might play an accessory role in mechanosensation; for example, they might amplify the mechanoreceptor potential, or modulate the primary mechanotransducer by mediating transient changes in calcium or other messengers (Vanden Abeele 2006). In the future, physiological characterization of UNC-7 hemichannel properties may distinguish between these hypotheses.

Although these results are the first to implicate UNC-7 as a mechanosensory molecule, other results are consistent with a role for innexins in *C. elegans* touch sensing. For example, it was shown recently (Sangaletti et al., 2014) that the TRNs express a mechanically gated current with innexin-like physiological and pharmacological properties. However, the mechanically gated currents that they identified are intact in *unc-7(e5)* animals, indicating that UNC-7 is not an essential component of this particular current (R. Sangaletti and L. Bianchi, personal communication). It is unclear how the pneumatic pressure stimulus used in these studies relates to externally-applied gentle touch, and one possibility is that different mechanically sensitive innexins function over different sensitivity ranges.

### UNC-7 and MEC-4 function independently in touch neurons

Unexpectedly, we have found that two mechanosensory channels, UNC-7 and MEC-4, are both required for normal touch responses in the TRNs; loss-of-function of either UNC-7 or MEC-4 alone leads to significant touch insensitivity. Nonetheless, several lines of evidence suggest that UNC-7 and MEC-4 function independently as mechanosensors, rather than functioning together in a common mechanotansduction complex. First, although both UNC-7 and MEC-4 proteins are distributed in a punctate pattern along the TRN dendrite, UNC-7- and MEC-4-containing puncta do not colocalize, and therefore appear to represent physically-distinct complexes. Second, although *unc-7* and *mec-4* single mutants both are insensitive to gentle touch, overexpression of *mec-4* can suppress the *unc-7* defect, while *unc-7* overexpression can partially suppress the gentle touch defect of *mec-4*. Third, *mec-4* and *unc-7* mutations have distinct phenotypes with respect to harsh touch responses in the TRNs; *unc-7* affects and is required for large, long-lasting *mec-4* independent responses, whereas *mec-4* is required for small, transient responses that resemble responses to gentle touch. Thus, although the activities of both UNC-7 and MEC-4 appear to be essential for sensitivity to weaker stimuli, either UNC-7 or MEC-4 is sufficient on its own to respond to stronger stimuli such as harsh touch.

Why would the TRNs need two or more mechanosensors? One possibility is that UNC-7 and MEC-4 respond to qualitatively distinct types of mechanical stimulation. MEC-4, for example, is believed to be tethered to both the extracellular matrix and the cytoskeleton (Arnadottir and Chalfie, 2010), whereas UNC-7, like the bacterial Msc, might directly sensing membrane tension via lipid interactions, (Kung et al., 2010). These different force detection mechanisms might in principle confer distinct biomechanical properties, resulting in specificity in the precise mechanical forces to which they respond. If neuronal response to light stimuli requires coincident activation of both UNC-7 and MEC-4, due to summing of these distinct inputs, this might also serve to filter out noise and improve the fidelity of response to small but behaviourally significant stimuli. In this context, it is interesting to note the recent demonstration (Rocio Servin-Vences et al., 2017) that both TRPV4 and PIEZO are required for mechanosensation in chondrocytes, and that they appear to function in distinct ways. PIEZO responds to membrane stretch, while TRPV4 appears to rely on tensile forces transmitted via the matrix. In addition, mechanosensory responses are known to be subject to modification by neuromodulators such as dopamine; in principle MEC-4, UNC-7, or both mechanosensory complexes may be targets for this modulation.

### UNC-7 plays genetically-distinct roles in gap junctions and mechanosensory hemichannels

In addition to its direct role in mechanosensation, UNC-7, along with UNC-9, also contributes to gap junctions that functionally link the anterior TRNs. In the locomotion circuit, for example in the electrical synapses between the AVB premotor interneurons and the B-class motorneurons, UNC-7 and UNC-9-containing gap junctions appear to be asymmetric, with UNC-7 expressed in AVB and UNC-9 expressed in B motor neurons (Starich et al., 2009). Likewise, UNC-7 and UNC-9 also form heterotypic gap junctions between AVA and A motor neurons (Kawano et al., 2011). In contrast, in the touch neurons, both innexins have been reported to be expressed in both AVM and ALM (Altun et al., 2009; Starich et al., 2009), and the *unc-7* mechanosensory phenotypes likewise imply expression in both ALM and AVM. Thus, for the ALM-AVM gap junctions, it seems likely that UNC-7 and UNC-9 contribute in both partner cells. Unlike UNC-7, the role of UNC-9 appears to be confined to gap junctions, since *unc-9* mutations (like AVM ablations) only affect responses to contralateral stimuli.

Although UNC-7 appears to function in both gap junctions and mechanosensory hemichannels, these two functions are genetically separable. For example, we have shown that whereas the L isoform of UNC-7 can rescue the mechanosensory defect of *unc-7(e5)*, the shorter SR isoform fails to rescue. Conversely, others (Starich et al., 2009) have shown that UNC-7SR and S can restore gap junction activity in the locomotion circuit, whereas UNC-7L could not rescue gap junction-dependent behavioural phenotypes and that UNC-7L could not produce gap junction currents in vitro. Since UNC-7L has been shown to form hemichannels when expressed in neuro2A cells (Bouhours et al., 2011), this suggests a distinction between isoforms, with UNC-7SR and S being required for gap junctions, while UNC-7L is capable of forming mechanosensitive hemichannels. The three isoforms differ only in the length of the N-terminal cytoplasmic region (see (Starich et al., 2009) for full details). The N-terminal region that is unique to UNC-7L is rich in proline, indicating a potential role in interaction with WW domain-containing proteins (Kay et al., 2000), and several other amino acid repeats that are suggestive of protein interaction motifs. Thus, this extended domain might mediate differential localisation or trafficking of UNC-7, or alternatively could interfere, either directly or through inter-protein interactions, with assembly into gap junctions. If UNC-7L is indeed the only isoform to be mechanosensitive, this unique domain could also hold the key to understanding the molecular basis of force sensing.

### Functional conservation of UNC-7 with mammalian pannexins

Intriguingly, we observed that the mechanosensory function of *unc-7* could be complemented by a mammalian homologue, the *Panx1* pannexin gene. Expression of a mouse *Panx1* transgene fully rescued the touch-insensitive defect of an *unc-7* null mutant, indicating strong functional conservation across a large phylogenetic distance. Although pannexins have not been directly implicated in touch or other somatosensory processes in vertebrates, Panx1 has been shown to be mechanically sensitive, mediating the release of ATP in a variety of cell types in response to membrane stretch. Our finding of a role for UNC-7, and its functional complementation by Panx1, suggests the possibility that pannexins might play undiscovered roles in touch or other mechanical senses in vertebrates.

Pannexins have been implicated in a huge array of medical conditions, including ischaemia-induced seizure, inflammation, hypertension, tumour formation and metastasis, and neuropathic pain, and are thus an important target for therapeutic intervention. A significant obstacle is their involvement in so many functions, requiring a deep understanding of these roles in order to intervene specifically. However, even their basic properties (non-selective versus anion-selective; high conductance versus low conductance) remain controversial (Chiu et al., 2014; Good et al., 2015). Our observation that mouse pannexin 1 can be functionally expressed in *C. elegans* neurons opens the door to a tractable model organism in which to study pannexin itself. As UNC-7 fulfils multiple functions in different cell types, understanding how this is determined will provide clues as to how this is achieved for pannexins in higher organisms.

## Experimental Procedures

### *C. elegans strains*. Strains used in this study are described in Table S1

*Plasmid constructs. Pmec-7*∷dsRNA plasmids for *unc-7, unc-9, inx-7* were constructed by ligating a cDNA fragment of approximately 600bp between the second and third multiple cloning sites of pPD117.01 (A gift from Andrew Fire). An *E. coli Cat1* (chloramphenicol acetyl transferase gene) dsRNA plasmid was constructed in a similar fashion. For each target, two plasmids, with sense and antisense orientations of the insert, were co-injected at 50ng/µl each. *unc-7* rescue plasmids were made using the Multisite Gateway^®^ 3-Fragment Vector Construction Kit (Invitrogen). A 1078bp *mec-4* promoter fragment (as previously used, Suzuki et al. 2003), was cloned into pDONR^™^ P4-P1R. *unc-7* (isoforms a and c) cDNA were amplified from RB1 cDNA library (a gift from Robert Barstead) and cloned into pDONR^™^ 221. These were combined with pDONR^™^ P2R-P3/SL2∷mcherry (a gift from Mario de Bono) in a derivative of pDEST^™^ R4-R3 into which *unc-54* 3’UTR had been inserted downstream of the recombination sites. Cysteine to arginine substitutions were made in the relevant pDONR^™^ 221 plasmid, using codon-optimised mutagenic primers, designed using *C. elegans* Codon Adapter (http://worm-srv3.mpi-cbg.de/codons/cgi-bin/optimize.py, (Redemann et al., 2011)). Partially overlapping complementary mutagenic primers were used to amplify the plasmid using Phusion^®^ High-Fidelity DNA Polymerase (Thermo Scientific), then *Dpn*I digestion was used to remove bacterially-derived template DNA, before transformation into *E. coli*. These were assembled into the *Pmec-4* expression vector in the same way as the wild type sequences. Pannexin 1 and 2 were amplified from a mouse cDNA library, and assembled into the *Pmec-4* expression vector in the same way. PVD and ASK expression vectors were constructed using the same Gateway^®^ strategy, using *ser-2prom3* and *sra-9* promoters, respectively (gifts from Marios Chatzigeorgiou and Lorenz Fenk (Chatzigeorgiou et al., 2010; Fenk and de Bono, 2015). The plasmids were injected at 50ng/µl. TRN-specific *mec-4*∷mcherry and *unc-7a*∷gfp fusions were constructed in a similar way, using the *mec-4* promoter, and injected at 20ng/µl and 50ng/µl, respectively. A second *unc-7a*∷gfp encoded a fusion, where GFP was inserted in the internal loop, between N290 and I291, surrounded by Gly Gly linkers, exhibited the same localisation.

Ca^2+^ *imaging.* Ca^2+^ imaging of anterior body touch stimulation of glued animals was essentially as described previously (Kerr et al., 2000; Suzuki et al., 2003), using a 1 second “buzz” stimulus just posterior of the terminal bulb. Posterior harsh body touch (for PVD stimulation) and nose “buzz” stimulation (for ASK stimulation) were performed as described previously (Chatzigeorgiou et al., 2010; Kindt et al., 2007). Images were recorded at 10 Hz using an iXon EM camera (Andor Technology), captured using IQ1.9 software (Andor Technology) and analysed using a custom written Matlab (MathWorks) program (Rabinowitch et al., 2013). For thermosensation, animals were glued in the usual way then treated by perfusion of buffer at the temperatures indicated. Mechanical stimulation for TRN and PVD imaging was performed in Neuronal Buffer (145mM NaCl, 5mM KCl, 5mM CaCL_2_, 5mM MgCl_2_, 20mM glucose, 10mM HEPES, pH7.2). Thermal stimulation and nose touch stimulation were performed in CTX (25mM KPO_4_ pH6, 1mM CaCl_2_, 1mM MgSO_4_). Where appropriate, neurons were categorised into ipsilateral or contralateral, depending on the position of their cell body with respect to the site of stimulation and a hypothetical midline.

### Channelrhodopsin experiments

L4 animals were transferred to retinal plates (made by seeding 55mm NGM plates with 160μl of a 1000:4 mixture of OP50 culture and 100mM all-*trans* retinal in ethanol and incubating overnight at 22°C), and grown at 22°C, then assayed as day 1 adults. Assay plates were prepared by seeding 30mm NGM plates with 40µl of a 1000:1 mixture of OP50 culture and 100mM all-*trans* retinal in ethanol and incubating for 30 minutes at 22°C. Control plates contained ethanol without all-*trans* retinal. Animals were picked individually to assay plates using an eyelash hair, and acclimatised for 15 minutes. To stimulate the TRNs, *lite-1(ce319)* worms expressing *Pmec-4∷ChR2* (Rabinowitch et al., 2016) were illuminated for 1 second with 1mW/mm^2^ blue light using 470nm LEDS controlled by a LEGO Mindstorms Intelligent NXT Brick.

### Laser killing of AVM

Laser ablation was carried out in L1/L2 animals as described by (Bargmann and Avery, 1995).

### Behavioural assays

Gentle and harsh body touch were performed on day 1 adults, by stroking with an eyelash hair or prodding with a platinum wire pick, respectively (Chalfie et al., 2014) just behind the pharynx terminal bulb (a3, **figure 1A**), except where stated. For habituation, stimulation was applied every 10 seconds, or as soon as possible thereafter if not moving forwards.

### Confocal microscopy

Images were acquired using a Zeiss LSM 780. Colocalisation was visualised using the Colocalization Finder plugin (C. Laummonerie and J. Mutterer, Strasbourg, France) for ImageJ. Object-based colocalisation of puncta was analysed using the JACoP plugin (Bolte and Cordelieres, 2006) for ImageJ, using particle centre of mass coincidence.

## Author contributions

Conceptualisation, D.S.W and W.R.S; Methodology, D.S.W and W.R.S; Investigation, D.S.W; Formal Analysis, D.S.W.; Writing – Original Draft, D.S.W.; Writing – Review & Editing, D.S.W and W.R.S.; Funding Acquisition, W.R.S.

## Acknowledgements

We are very grateful to Yee Lian Chew, Kristin Webster and Yiquan Tang for critical reading of the manuscript, and members of the Schafer, de Bono and Taylor labs for helpful discussions. We are grateful to the LMB workshops for help with the channelrhodopsin setup. We thank Andrew Fire, Robert Barstead, Lorenz Fenk, Mario de Bono and Marios Chatzigeorgiou for plasmids and strains. Some strains were provided by the CGC, which is funded by NIH Office of Research Infrastructure Programs (P40 OD010440). This work was funded by the Medical Research Council (MC_A023_5PB91) and Wellcome Trust (WT103784MA).

## Competing interests

No competing interests declared.

**Table S1.**
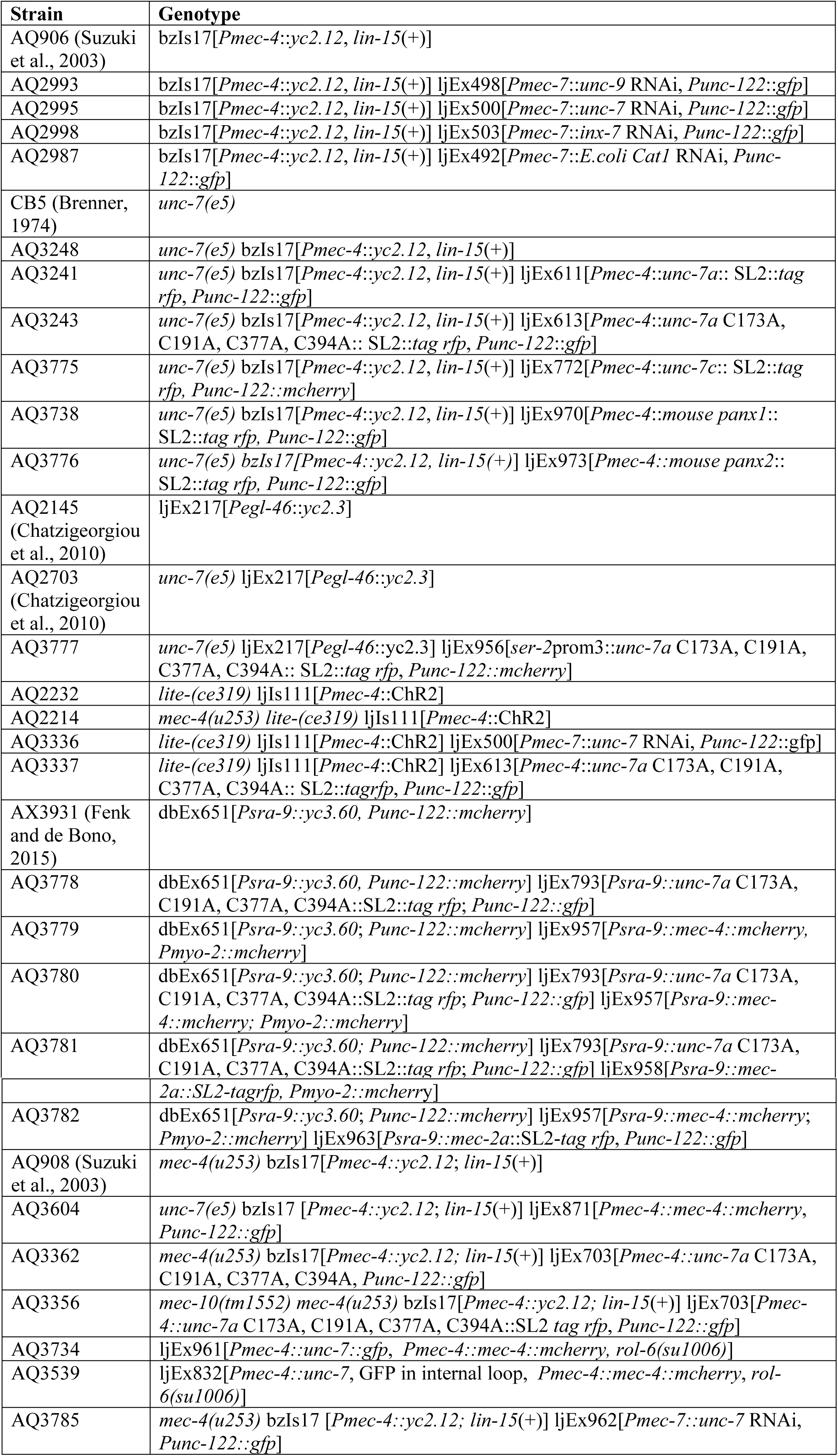
Strains used in this study.

